# Complementary prefrontal and thalamic representational dynamics during response planning

**DOI:** 10.64898/2026.09.07.749911

**Authors:** Daniel Hähnke, Ajit Ranganath, Xuanyu Wang, Salem Ayasreh Fierro, Tobias W. Bernklau, Simon N. Jacob

## Abstract

Advance contextual information can guide behavioral responses, but how prospective action representations unfold across prefrontal and thalamic neuronal populations of the cognitive control network remains unclear. Here, we trained mice on a contextual response-planning task in which auditory cues either predicted the upcoming instructed movement or left response identity unresolved until a later instruction. Predictive contexts improved accuracy and produced subthreshold, directionally biased movements, indicating that mice used contextual information before instruction onset. Extracellular recordings revealed prospective response-side information in prelimbic cortex (PL) and mediodorsal thalamus (MD), but with distinct dynamics. MD represented prospective response side earlier during the context epoch and showed a planning-related temporal advance after instruction onset. In contrast, PL exhibited enhanced instruction- epoch coding and stronger cross-epoch generalization, which could be localized to sparse neuronal subpopulations. These complementary representational dynamics suggest distinct but coordinated roles for prefrontal-thalamic circuits in using advance information to support flexible action planning.

## Introduction

Contextual information can be used to plan an upcoming action before the final imperative instruction appears (Churchland, 2024; Mukherjee et al., 2021). However, when the available information is insufficient, a premature commitment must be withheld until a later cue resolves the appropriate response (Carlsen & MacKinnon, 2010). Understanding how neuronal populations represent an upcoming response under these two conditions is important for explaining how behavior benefits from advance information. More broadly, context-dependent use or withholding of advance preparation can be viewed as one aspect of cognitive flexibility (Hanganu-Opatz et al., 2023), commonly operationalized as selecting an appropriate response, rule or stimulus-response mapping under changing task demands (Kaefer et al., 2020; Miller & Cohen, 2001; Wimmer et al., 2015).

The prefrontal cortex has long been implicated in cognitive flexibility and executive control (Hanganu-Opatz et al., 2023; Miller & Cohen, 2001). Electrophysiological studies in non-human primates and humans show that lateral prefrontal neurons represent abstract rules, task sets, goals and decision variables during flexible behavior (Mansouri et al., 2006; Mian et al., 2014; Wallis & Miller, 2003). In rodents, related functions have been attributed primarily to medial prefrontal cortex, particularly the prelimbic area (PL), which has been associated with rule switching, context-dependent response selection and maintenance of task-relevant information across delays (Kaefer et al., 2020; Nakayama et al., 2018; Spellman et al., 2021). Importantly, prefrontal representations of higher-order behavioral and motivational states have been shown to fully decouple from lower-level sensory or motor representations (Rautio et al., 2026), arguing that this region may support context-dependent cognitive planning, rather than motor response preparation.

Prefrontal cortex is reciprocally connected with the mediodorsal thalamus (MD), which has emerged as an active contributor to cognitive flexibility in its own right (Halassa & Kastner, 2017; Parnaudeau et al., 2015). In rodents, MD perturbation impairs flexible updating of stimulus-outcome and cue-context associations, including reversal learning and rule switching (Lam et al., 2024; Parnaudeau et al., 2013; Parnaudeau et al., 2015; Rikhye et al., 2018). Evidence from macaques similarly implicates MD in adaptive learning and flexible choice when task contingencies change (Chakraborty et al., 2016). Experimental studies further indicate that MD inputs can amplify or sustain prefrontal activity and influence transitions between task-relevant cortical states (Rikhye et al., 2018; Schmitt et al., 2017).

Together, the available literature identifies PL and MD as interacting components of a distributed cognitive control system. However, their respective contributions, and thus the division of labor between prefrontal and thalamic populations in planning upcoming actions, remain unclear. Classical delayed-response tasks generally provide decisive, deterministic information about the future response and examine its representation during a memory delay (Curtis et al., 2004; Funahashi & Andreau, 2013; Fuster & Alexander, 1973), whereas cognitive-flexibility tasks often emphasize switches between rules (Bolkan et al., 2017; Parnaudeau et al., 2013), sensory modalities (Lam et al., 2024; Rikhye et al., 2018; Schmitt et al., 2017; Wimmer et al., 2015) or action-outcome associations (Bernklau et al., 2024; Tafazoli et al., 2026). These tasks have clarified how prefrontal and thalamic populations maintain or update task representations, but they do not directly compare neuronal dynamics when advance response planning is possible with those observed when response identity remains unresolved.

Here, we addressed this gap using a contextual response planning task in which mice first received an auditory context cue that was either predictive or non-predictive of the upcoming instruction. Predictive contexts thus specified the forthcoming required response, a movement to the left or right side, whereas non-predictive contexts left response identity unresolved until the instruction appeared. Because context predictiveness varied across trials, the task contrasted advance planning with deferred response specification within the same behavioral sequence. Using high-density extracellular recordings from PL and MD, we examined how population information about response side emerged and evolved across the context, instruction and execution epochs. In both regions, population activity distinguished the prospective response side after predictive cues and subsequently expressed a context-invariant instructed-side representation after instruction onset. MD showed earlier prospective side decoding during the context epoch and a planning-related temporal advancement during the instruction epoch. PL, in contrast, showed a planning-related magnitude enhancement during the instruction epoch and stronger cross-epoch generalization concentrated in sparse neuronal subpopulations. Our findings of complementary representational dynamics regarding the timing and stability of action representations in the rodent prefrontal-thalamic circuit provide a population-level account of context-dependent action planning with broader implications for cognitive flexibility.

## Results

### Predictive context cues support response planning

We trained head-fixed mice (n = 5) on an auditory contextual response planning task in which predictive response information was available on some trials but not others. The task consisted of a context epoch followed by an instruction epoch (**Fig. 1a**). After trial initiation, one of three auditory stimuli (context cues) was presented for 100 ms: left-plan (low-pass filtered white noise, predicting a go-left instruction), right- plan (high-pass filtered white noise, predicting a go-right instruction) or defer (unfiltered white noise, non-predictive, with a 50 % probability of a subsequent go-left or go-right instruction). After a delay (minimum 500 ms; see below), a second auditory stimulus (instruction cue) indicated the correct response side (ascending-pitch sound for go-left and descending-pitch sound for go-right). Based on the context- instruction pairing, we defined four trial types. Trials with predictive context were left-plan (Lp) and right- plan (Rp), whereas trials with non-predictive context were left-defer (Ld) and right-defer (Rd). Mice reported the instructed side by rotating a response ball (Sanders & Kepecs, 2012) after instruction onset. A response was registered when ball rotation exceeded a fixed threshold (57.6 ° or 8 mm). The instruction cue was presented for up to 1000 ms and was terminated earlier upon threshold crossing. Correct responses were rewarded with a drop of water, whereas trials without a report within the permitted response window (up to 5000 ms after the instruction cue) were classified as misses. The animals were trained to keep the ball stationary during the delay following the context cue. If they rotated the ball beyond a small threshold (18 ° or 2.5 mm) during the delay, the timer was reset.

**Fig. 1.**
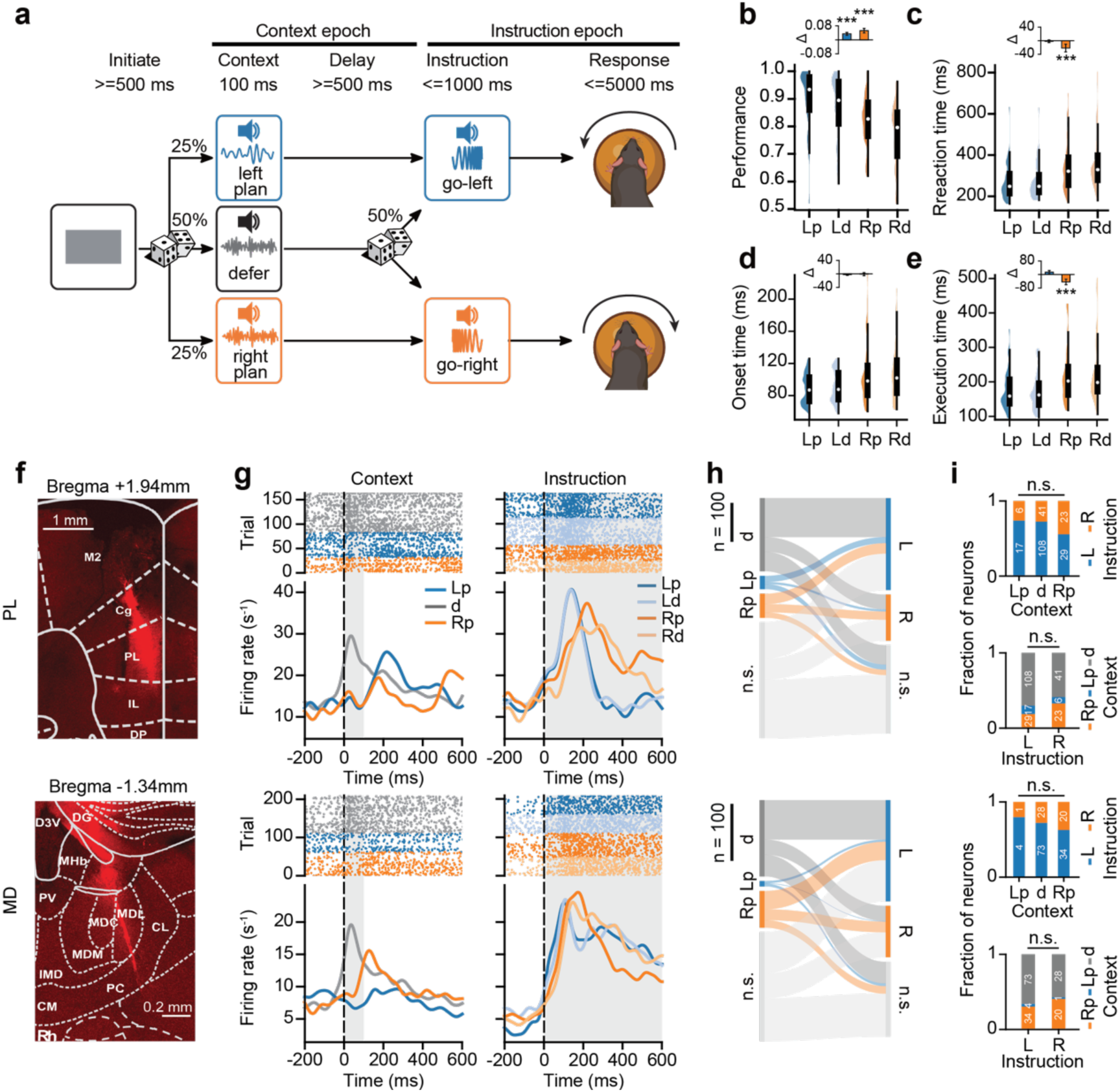
Contextual planning task and extracellular neuronal recordings. **a.** Schematic of task design. **b.** Task performance in all four conditions. Distributions across sessions are shown as half-violins with box plots indicating the median (white dot), quartiles (thick line), and 1.5 times the interquartile range (thin line). Lp, left-plan; Ld, left-defer; Rp, right-plan; Rd, right-defer. Inset: context effect for left or right instruction side. Asterisks indicate significance of Wilcoxon signed-rank tests: \**p* < 0.05; \*\*\**p* < 0.001. **c.** Reaction time in all four conditions, defined as the interval from instruction onset to response-threshold crossing. **d.** Response-onset time, defined as the interval from instruction onset to the first extracted submovement. **e.** Response-execution time, defined as the interval from response onset to threshold crossing. **f.** Example coronal brain sections showing silicon-probe tracks labeled with DiI in prelimbic cortex (PL, top) and mediodorsal thalamus (MD, bottom), overlaid on the Allen Brain Atlas (Wang et al., 2020). **g.** Responses of example PL (top) and MD (bottom) neurons after context (left) and instruction (right) cues. Raster plots are displayed above mean firing rates, separated by context or instruction conditions. **h.** Transitions in neuronal selectivity between the context epoch (left column; d, defer) and the instruction epoch (right column) in PL (top) and MD (bottom). Bar height is proportional to the number of selective units. n.s., not significant. **i.** Distribution of instruction-cue selectivity as a function of context-cue selectivity, and vice versa, for PL (top) and MD (bottom).

To assess whether mice utilized the predictive context cue, we first analyzed behavioral performance. Accuracy was significantly higher in plan trials than in defer trials (**Fig. 1b**; repeated-measures two-way ANOVA, main effect of context: *F*(1,52) = 29.5, *p* < 0.001). Although all animals exhibited a response bias toward the left side (main effect of instruction: *F*(1,52) = 21.2, *p* < 0.001), the accuracy improvement in plan relative to defer trials was significant for both response directions (**Fig. 1b**, inset). Reaction time, defined as the interval between instruction onset and threshold crossing of the response ball, showed a marginal context effect (**Fig. 1c**; repeated-measures two-way ANOVA, main effect of context: *F*(1,52) = 3.7, *p* = 0.061). A significant reaction time advantage for plan compared to defer trials was observed only for go-right trials, but not for go-left trials (**Fig. 1c**, inset).

Although animals were trained to withhold overt movement responses during the delay, the response ball revealed subthreshold behavioral fluctuations below the report-detection threshold. To quantify these fluctuations, we analyzed the continuous rotational readout of the response ball by decomposing its velocity traces into minimum-jerk submovements (SMs) using an iterative fitting algorithm with customized bell-shaped kernels (**Fig. S1a**; see Methods). This approach provided estimates of the magnitude (peak velocity), duration and direction of individual SMs. Following predictive context cues, accumulated SMs during the delay tended toward the predicted side (**Fig. S1b**, top and bottom; quantified using a side index based on fitted SM parameters, **Fig. S1c-e**). Defer trials showed even stronger and more prolonged SM activity (**Fig. S1b**, middle), predominantly biased toward the left side (**Fig. S1c-e**), consistent with the behavioral bias observed in accuracy (**Fig. 1b**) and reaction time (**Fig. 1c**). Thus, leaving response identity unresolved did not eliminate subthreshold motor activity. During the instruction epoch, overt reports differed markedly from the subthreshold SMs observed during the delay. Report-related SMs were up to ten times larger in magnitude (**Fig. S1f**) and showed no significant differences between plan and defer contexts (**Fig. S1g-i**; no main effect of context on count, peak velocity or duration).

We analyzed leftward and rightward SMs separately and found that context-dependent SM directionality was mainly driven by rightward SMs: their count (repeated-measures ANOVA, *F*(2,98) = 35.7, *p* < 0.001), peak velocity (repeated-measures ANOVA, *F*(2,98) = 35.7, *p* < 0.001) and duration (repeated-measures ANOVA, *F*(2,98) = 7.3, *p* < 0.01) increased with the likelihood of an upcoming go-right instruction, whereas leftward SM count and magnitude remained largely stable across contexts (**Fig. S1j-l**). During the instruction epoch, SMs were directed toward the instructed side, and prior context no longer modulated SM count, peak velocity or duration (**Fig. S1m-o**).

The reaction time difference between right-plan and right-defer trials (**Fig. 1c** inset) could arise either from earlier response initiation or from faster response execution. To distinguish these possibilities, we separated reaction time into response onset time (the interval from instruction onset to the start of the first SM) and response execution time (the interval from the first SM to threshold crossing). No significant context-related differences were observed in the response onset time (**Fig. 1d**). However, a context-related difference was present in response execution time for go-right trials (**Fig. 1e**).

In summary, mice used predictive context cues before the instruction: predictive contexts were associated with higher accuracy, and go-right trials also showed a reaction-time benefit that reflected shorter response execution. Animals exhibited subthreshold behavioral fluctuations after both predictive and non-predictive context cues. Thus, the defer condition was not a state without movement planning; instead, the non-predictive cue left response identity unresolved, whereas predictive cues biased subthreshold movements toward the upcoming instructed side.

### PL and MD single units show distinct context- and instruction-related dynamics

Having established that mice used predictive context information before the instruction, we next sought to characterize neuronal activity in PL and MD, two interconnected hubs of the brain’s cognitive control system. First, we confirmed the anatomical relationship between these regions. In two separate experiments, we injected cholera toxin subunit B (CTB) into either PL (**Fig. S2a**) or MD (**Fig. S2c**). CTB is taken up by axon terminals near the injection site and transported retrogradely to the corresponding neuronal cell bodies, thereby labeling neurons that project to the injected region. We divided PL along the anterior-posterior axis into anterior (aPL), middle (mPL), and posterior (pPL) subregions, and MD along the medial-lateral axis into medial MD (MDm) and lateral MD (MDl). MD-to-PL projections followed a spatial gradient, with MDm neurons projecting preferentially to aPL and MDl neurons to pPL (**Fig. S2b, e, f**). PL-to-MD projections reached in particular the MDm, whereas more superficial regions in the medial prefrontal cortex (e.g., cingulate and secondary motor cortex) connected predominately to the MDl (**Fig. S2d, g, h**). These results confirmed reciprocal PL-MD connectivity (Collins et al., 2018; Krettek & Price, 1977) and indicated a topographic organization along the anterior-posterior axis in PL and the medial-lateral axis in MD.

Next, we recorded prefrontal and thalamic single-unit activity using 32-channel silicon probes (NeuroNexus) while mice performed the planning task (**Fig. 1f**). Spike waveforms were sorted using Kilosort1 (Pachitariu et al., 2016), followed by manual curation. We recorded 1020 units in PL and 711 in MD across 38 sessions from four animals. Of these, 629 PL and 420 MD units met the inclusion criteria (curated single units with a firing rate above 1 spike/s and at least 5 correct trials per condition) and entered subsequent analyses.

Single units in both PL and MD responded to context and instruction cues, typically showing rapid firing increases at stimulus onset followed by sustained activity during the delay and instruction epochs (**Fig. 1g**). We used the parameter-free ZETA (ζ) test (Montijn et al., 2021), which compares the cumulative distribution of spike times with a shuffled baseline, to identify cue-responsive units (**Fig. 1h**; context cue: 48.8 % of PL units and 52.1 % of MD units; instruction cue: 60.9 % of PL units and 67.1 % of MD units). The maximal ζ value defined the strength and timing of a unit’s response. For each significantly responsive unit, we assigned cue preference as the context or instruction cue yielding the maximum ζ within a 600 ms window after context onset or a 300 ms window after instruction onset. In both regions, more units preferred the defer context cue than the plan cues, and among the plan cues, more preferred Rp than Lp (**Fig. 1h**, left). By contrast, during the instruction epoch, more units preferred the go-left than the go-right instruction (**Fig. 1h**, right). No statistically significant association was detected between context-cue and instruction- cue preferences (**Fig. 1i**; PL: contingency test χ2(3, N = 383) = 6.29, p = 0.098; MD: χ2(2, N = 307) = 8.50, p = 0.075).

To capture the collective dynamics of PL and MD, we examined population-averaged firing rates from condition-balanced correct trials across sessions. Units in both regions exhibited an early peak (∼50 ms) following the defer context and later peaks (∼200 ms in PL and ∼100 ms in MD) following the plan contexts (**Fig. 2a, b**, left), with Rp eliciting stronger activity than Lp. In PL, the firing-rate difference between Lp and Rp declined after this initial peak and was no longer evident before instruction onset, whereas in MD the difference persisted throughout the context delay (**Fig. 2a, b**, right). Across both regions, go-left instructions elicited stronger neuronal activity than go-right instructions. Notably, during the first 100 ms after instruction onset, PL units showed a significant difference between plan and defer contexts (Wilcoxon signed-rank test: Lp vs. Ld, p < 0.001; Rp vs. Rd, p = 0.010); the difference was not significant in the subsequent 100 ms window (Lp vs. Ld, p = 0.090; Rp vs. Rd, p = 0.146). No comparable difference was detected in MD. Matching this pattern, PL units displayed a biphasic increase in firing during the instruction epoch: an initial rapid rise around 10 ms after instruction onset, and a second peak around 100 ms after instruction onset (**Fig. 2c**). In contrast, MD units showed a single peak in firing-rate increase around 45 ms after instruction onset, with neuronal recruitment distributed more evenly throughout the instruction epoch and peak timing falling between the two PL peaks (**Fig. 2d**).

**Fig. 2.**
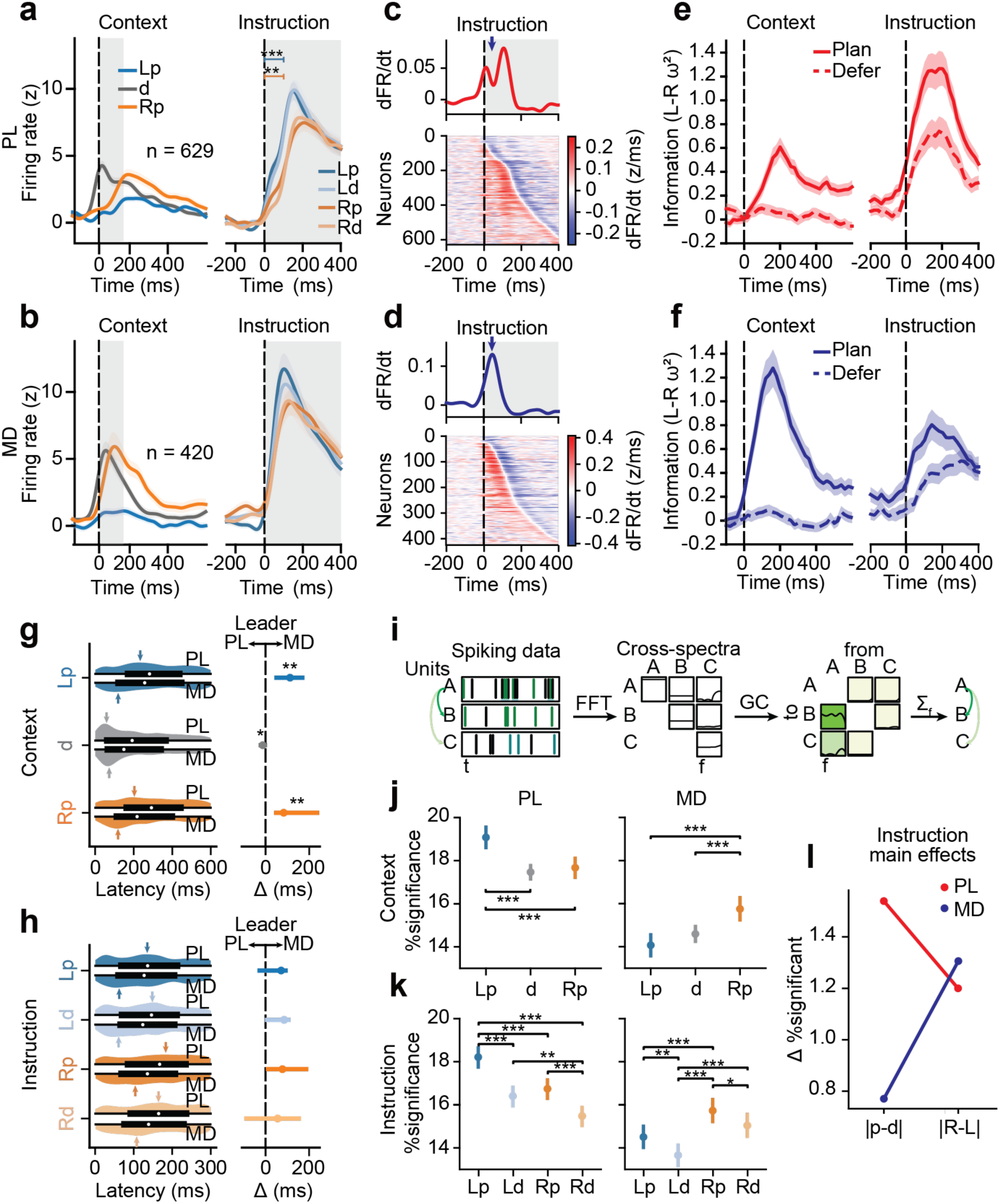
Neuronal responses to context and instruction cuesa. Mean normalized firing rates across PL units (n = 629), aligned to context onset (left) and instruction onset (right). Vertical dashed lines mark auditory-cue onset, and shaded areas mark cue presentation. Solid lines indicate the mean, shaded bands ± SEM. Significance marks: Wilcoxon signed-rank test; \*\**p* < 0.01, \*\*\**p* < 0.001. **b.** Same as **a** for MD neurons (n = 420). **c.** Change in firing rate after instruction onset, measured as the first temporal derivative of normalized firing rate (dFR/dt). Top, mean dFR/dt across PL neurons; bottom, dFR/dt across PL neurons sorted by peak-firing time after instruction onset. **d.** Same as **c** for MD. **e.** Information about the instructed response side (left vs. right) quantified as the percentage of explained variance (PEV) in PL firing rates, aligned to context onset (left) and instruction onset (right). Solid lines show the mean, while shaded bands representing ±SEM. **f.** Same as **e** for MD. **g.** ζ-test-derived single-unit response latency after the context cue. Left, distributions shown as half-violins with box plots indicate the median (white dot), quartiles (thick line), and 1.5 times the interquartile range (thin line); arrows mark the first mode of each kernel density estimate (KDE). Right, differences between the first KDE modes for PL and MD after each context cue. Dots indicate the measured differences and whiskers the bootstrapped 95% confidence interval. Bootstrap test: \**p* < 0.05, \*\**p* < 0.01. **h.** Same as **g** for the instruction cue. **i.** Spike-based Granger-Causality analysis using simulated Poisson spike trains. Binary spike trains from three simulated neurons (left; A influences B across frequencies and C in a high-frequency band) were Fourier transformed to obtain a spectral-density matrix (middle). Frequency-domain Granger-Geweke values were computed using Wilson’s matrix factorization, and values were averaged across frequencies to obtain a scalar measure of directed predictability (right). **j.** Percentage of intraregional unit pairs with significant spike-spike GC during the context epoch in PL (left) and MD (right). Significance for each pair was assessed using 200 trial shuffles. **k.** Same as **j** for the instruction epoch. **l.** Descriptive main effects of context (absolute difference between plan and defer trials, |p-d|) and instruction (absolute difference between left and right trials) on the percentage of significant intraregional GC pairs in PL and MD.

These region-specific and context-dependent firing patterns in PL and MD raised the question of how much firing-rate variability was associated with instructed response side. We therefore calculated the percentage of explained variance (PEV) in each unit’s firing rate attributable to instructed side (left vs. right) and compared it between plan and defer contexts. In both regions, PEV was significantly higher in plan than defer trials (**Fig. 2e, f**). Because the two predictive context cues were acoustically distinct and predicted different response sides, the context-epoch contrast contains both cue-identity and prospective response-side information. In the instruction epoch, however, these sensory effects were controlled for by the task design (the left/right instruction cues were the same for plan and defer), indicating stronger response-side information when the upcoming response had been specified in advance. In PL, the planning-induced PEV elevation was greater during the instruction epoch than during the context epoch, and vice versa in MD, revealing a regional dissociation in the relative prominence of instructed-side information across task epochs.

To compare the timing of cue-evoked neuronal responses in PL and MD, we examined the distributions of ζ peak latencies across recorded units under different task conditions. During the context epoch, the latency distributions were consistent with unimodality in both regions except for the Lp condition in MD (**Fig. 2g**, left; Hartigan’s dip test, Lp in MD: p = 0.021; all others p > 0.05). Following the defer cue, the first mode of the latency distribution, estimated using kernel density estimation (KDE), occurred earlier in PL than MD (**Fig. 2g**, right; PL = 59 ms, MD = 73 ms; permutation test, p < 0.05). For the predictive contexts, the first mode occurred earlier in MD than PL (Lp: PL = 234 ms, MD = 119 ms; Rp: P = 203 ms, MD = 119 ms; both p < 0.01). In the instruction epoch, PL latency distributions departed from unimodality in all but the right-plan condition (**Fig. 2h**, left; dip test for Rp: p = 0.305; all others p < 0.05), consistent with the biphasic dynamics in population firing rates. MD latency distributions were unimodal except for Rp (p = 0.045; all others p > 0.05), consistent with the single-peak population dynamics. Across conditions, the first MD modes were numerically earlier than those in PL, although these differences did not reach statistical significance (permutation test, **Fig. 2h**, right).

As an exploratory measure of within-region directed statistical dependence, we applied nonparametric, unconditional Granger-Geweke causality (GC) to spike trains from pairs of units within PL or within MD, using multitaper spectral estimation (Dhamala et al., 2008; Granger, 1969) (**Fig. 2i**). GC values were averaged across frequencies, and pairs exceeding a per-pair permutation threshold (p < 0.05; 200 trial shuffles) were classified as significant GC pairs. During the context epoch, both regions contained fewer significant GC pairs in defer than plan trials (**Fig. 2j**), despite higher mean firing rates after the defer cue (**Fig. 2a, b**). More significant pairs were observed for Lp in PL and for Rp in MD. During the instruction epoch, the fraction of significant pairs varied with context and instruction, with higher fractions in plan than defer trials and in go-left than go-right trials (**Fig. 2k**). The relative sizes of these effects also differed between regions, with contextual planning (plan vs. defer) dominating PL connectivity, and instructed side (left vs. right) dominating MD connectivity (**Fig. 2l**).

In summary, both PL and MD showed task-modulated activity, yet with complementary temporal dynamics and representational emphasis. PL neurons were more sensitive to whether contextual information was present or not, while MD neurons were more sensitive to the instructed response side. This dissociation was apparent in both the context and instruction epochs, and in both population activity and intraregional connectivity.

### Planning produces distinct shifts in PL and MD population codes

We next examined how information about upcoming left and right responses was represented at the population level. We trained support vector machine (SVM) classifiers on condition-balanced pseudo-population spiking activity pooled across all recorded units within each region.

First, we trained and tested decoders separately for plan and defer trials. In plan trials, left-plan and right- plan conditions were readily discriminated from both regions shortly after the context cue, reaching accuracy above 0.9, with MD peaking earlier than PL, in line with our previous results (**Fig. 3a, b**, top left; peak time after context onset: 240 ms for PL and 140 ms for MD; cp. **Fig. 2a, b**). After the instruction cue, instructed side was decoded rapidly from both regions with ceiling-level accuracy and comparable timing (**Fig. 3a, b**, top right; peak time after instruction onset: 162 ms for PL and 162 ms for MD). In defer trials, after instruction onset, both regions achieved accuracy above 0.9, with PL peaking earlier than MD (**Fig. 3c, d**, top right; peak time after instruction onset: 135 ms for PL and 203 ms for MD).

**Fig. 3.**
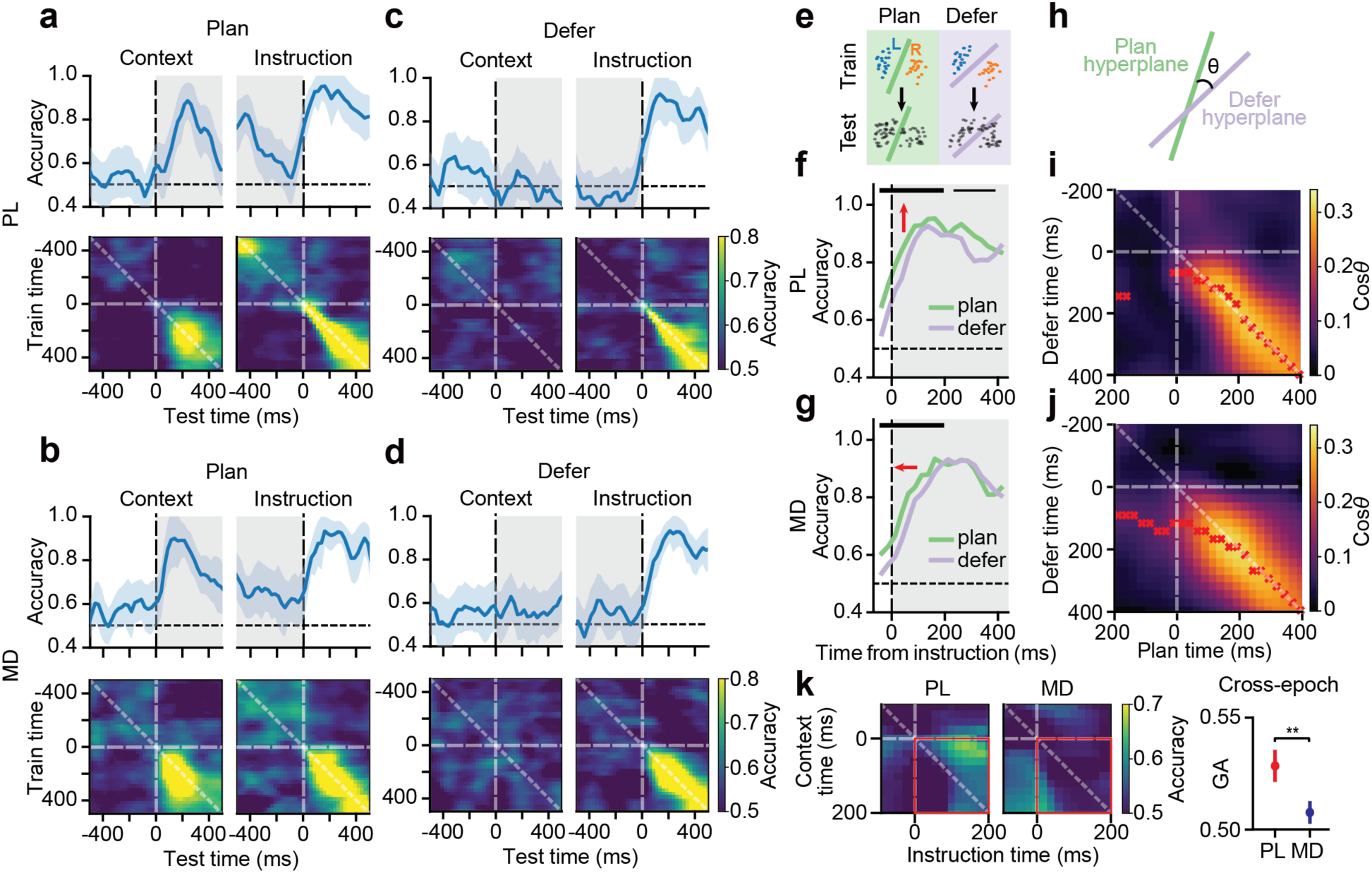
Population representations of instructed response side. **a.** SVM decoding accuracy for left-plan versus right-plan conditions (Lp vs. Rp) from PL population activity aligned to context-cue onset (left) and instruction-cue onset (right). PL units from all sessions were pooled across animals and sessions into a pseudo-population. Top, decoding accuracy at each time point from randomly selected, nonoverlapping training and test trials; lines and shading mark the mean and standard deviation across 100 shuffles. Bottom, cross-temporal decoding obtained by training at one time point and testing at another. Vertical dashed lines indicate cue onset, grey shading marks cue presentation, and the horizontal dashed line marks chance (0.5). **b.** Same as **a** for MD. **c.** SVM decoding accuracy for instructed side in defer trials (Ld vs. Rd) from PL activity aligned to context-cue onset (left) and instruction- cue onset (right). **d.** Same as **c** for MD. **e.** Within-context SVM decoding. Accuracy was determined by training and testing on randomly drawn, nonoverlapping sets of trials (n = 50) from the same context (plan or defer), with trial counts balanced between instructions. **f.** Instructed-side decoding accuracy in plan and defer trials in PL. Horizontal bars indicate significant time windows from permutation tests: thick bars, *p* < 0.01; thin bars, *p* < 0.05. **g.** Same as **f** for MD. **h.** Cosine similarity between SVM classifier hyperplanes. **i.** Temporally resolved cosine similarity between plan- and defer-context SVM hyperplanes for instructed- side decoding from PL pseudo-population activity. Horizontal and vertical dashed lines mark instruction onset, and red dots mark the maximum along each column. **j.** Same as **i** for MD. **k.** Cross-epoch generalization of response-side information in PL and MD. Right, generalization accuracy (GA), defined as mean cross-temporal decoding accuracy within the first 200 ms of each epoch (red rectangle in the left).

We then investigated the temporal stability of left-right representations using cross-temporal decoding (Meyers, 2018). In both regions, representations that discriminated future response side in plan trials appeared more stable during the context epoch than during the instruction epoch (**Fig. 3a, b**, bottom). In PL, instruction-epoch representations of instructed side appeared more stable in plan than defer trials (**Fig. 3a, c**, bottom right). In MD, instruction-epoch representations were comparable between contexts, with one stable pattern in the first 200 ms followed by another stable pattern thereafter (**Fig. 3b, d**, bottom right). Across contexts, PL showed more dynamic instruction-epoch representations than MD, in line with our previous findings of biphasic activity in the PFC (**Fig. 2a, b**).

To examine how planning modulated instructed-side representations, we trained and tested decoders separately for the two contexts (**Fig. 3e**). In PL, decoding accuracy was consistently higher in plan trials throughout the instruction epoch, producing an upward shift (**Fig. 3f**). Thus, advance information was associated primarily with a stronger instructed-side representation in PL. In MD, decoding was enhanced in plan trials only during the first 200 ms after instruction onset, producing a leftward shift (**Fig. 3g**). Thus, planning was associated primarily with earlier instructed-side information in MD.

Cross-context decoding between plan and defer trials in the instruction epoch (**Fig. S3a**) achieved accuracy comparable to within-context decoding, indicating similar instructed-side representations across contexts. A planning-related advantage was present in PL during the first 100 ms after instruction onset (**Fig. S3b**) but was absent in MD (**Fig. S3c**). Decoders trained on defer trials performed similarly in both contexts during the first 200 ms but became less reliable thereafter (**Fig. S3d, e**). We further compared the cosine similarity of SVM hyperplanes from plan and defer decoders over time (**Fig. 3h**). In PL, cross-context similarity peaked mainly along the diagonal (**Fig. 3i**), consistent with similar temporal dynamics and a planning-related magnitude shift (**Fig. 3f**). In MD, similarity peaks were off-diagonal during the first 200 ms after instruction onset (**Fig. 3j**), consistent with earlier instructed-side information in plan than defer trials (**Fig. 3g**).

Finally, we asked whether instructed-side information (Lp vs. Rp) generalized from the context epoch to the instruction epoch. We performed cross-epoch decoding using the first 200 ms of each epoch and defined mean cross-decoding accuracy in this window as the generalization accuracy (GA) (**Fig. 3k**). Because the predictive context and instruction cues were acoustically distinct but specified the same response side, successful cross-epoch decoding provides stronger evidence for a generalized response-side representation than context-epoch decoding alone. PL showed a significantly higher GA than MD, indicating more robust cross-epoch generalization in PL.

In summary, population decoding revealed largely context-general instructed-side representations during the instruction epoch in both regions. Advance planning was associated primarily with a magnitude enhancement in PL and a temporal advancement in MD. In addition, PL showed substantially stronger cross-epoch generalization than MD.

### Sparse PL populations support cross-epoch generalization

Although instructed-side information generalized across task epochs at the population level, single-neuron cue preferences were largely independent between epochs (**Fig. 1h, i**). This raised the question of how a generalizable population code could emerge from individual neurons whose preferences changed across epochs. Linear SVM decoding establishes whether instructed-side information is separable in the high-dimensional population state space and whether this separation generalizes across epochs, but it does not directly reveal how individual neurons contribute to the population dimensions supporting that generalization. Therefore, we next examined the structure of the population representation by decomposing task-related activity into interpretable dimensions and quantifying the contribution of individual neurons to each dimension. One possibility is that cross-epoch generalization depends disproportionately on a small, sparse subset of neurons, while task variable selectivity changes in the other neurons. In this case, traditional dimensionality reduction approaches that maximize orthogonal variance in the populational states space (e.g., Principle Component Analysis, PCA) may overlook the sparse neurons that contribute most strongly to the code (**Fig. 4a**).

**Fig. 4.**
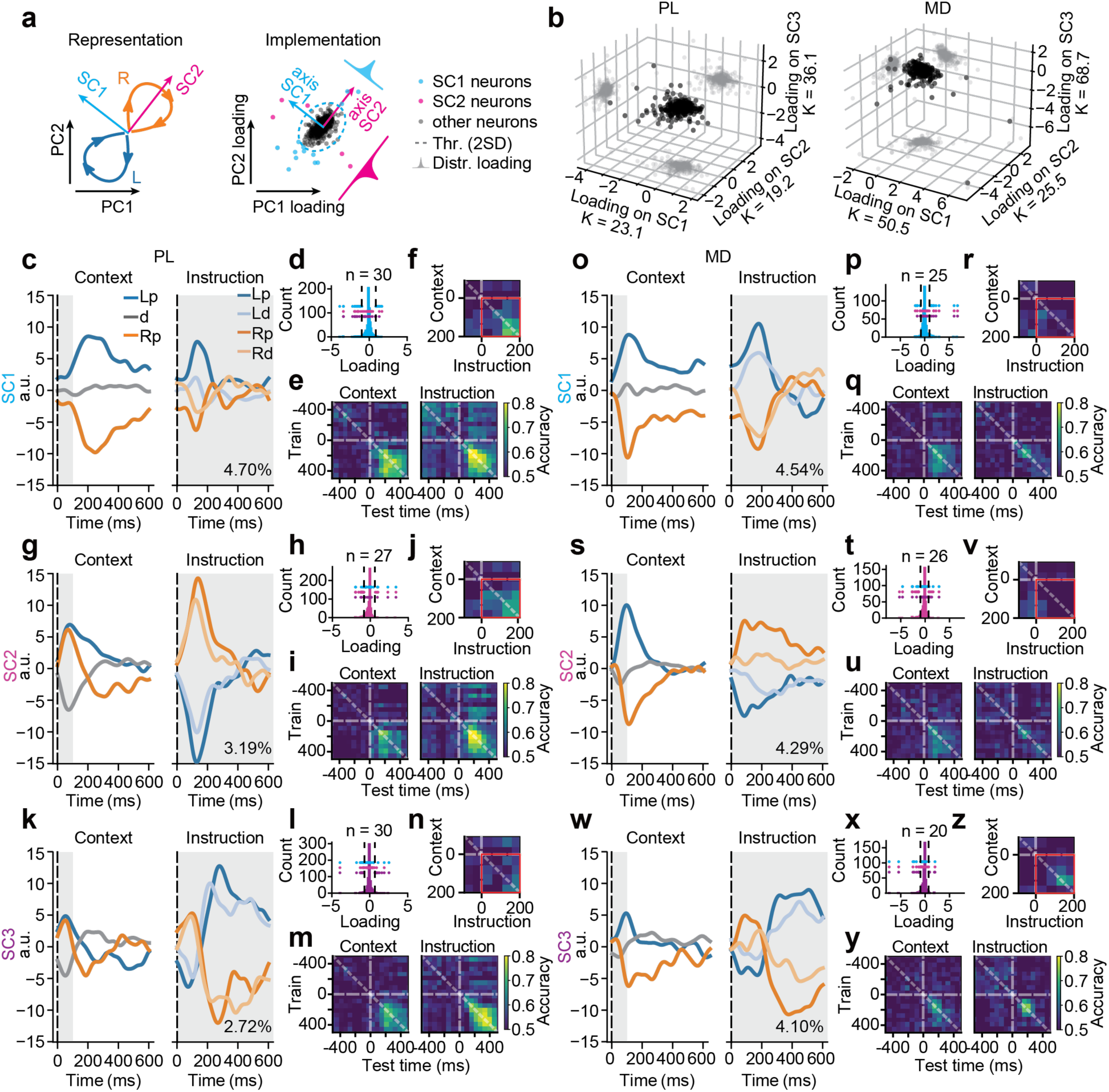
Sparse implementation of population representationsa. Multiple possible implementations of the same representational dynamics. Population activity in a high- dimensional neuronal state space can be projected onto principal components that maximize explained variance across time or onto alternative axes. Demixed sparse component analysis (dSCA) identifies projection axes that capture task-factor-related variance (left) while promoting sparse single-neuron loadings (right). **b.** Neuronal loadings onto the three identified sparse components (SCs) in PL (left) and MD (right). K, kurtosis. **c.** Activity of PL sparse component 1 (SC1; 4.70% variance explained) across task conditions during the context and instruction epochs. SCs were identified using plan trials and then applied to all conditions. **d.** Distribution of neuronal loadings onto SC1. Dominant neurons for each SC (absolute loading above 2 SD threshold) are overlaid on the full SC1 loading distribution. Dashed lines mark the threshold for dominant-neuron classification. **e.** Cross-temporal decoding accuracy for left-right task information from SC1-dominant neurons during the context (left) and instruction (right) epochs. **f.** Cross- epoch generalization within the first 200 ms of each epoch. **g-j.** Same as **c-f** for PL SC2. **k-n.** Same as **c-f** for PL SC3. **o-z.** Same as **c-n** for MD sparse components (dominant neuron threshold 1.5 SD).

To identify possible sparse neuronal populations, we used demixed sparse component analysis (dSCA), previously developed in our laboratory (Lin et al., 2023). We first demixed population activity to isolate variance associated with left-right task conditions from condition-independent activity, including global temporal dynamics and sensory-evoked responses (Kobak et al., 2016). Unlike PCA, which identifies orthogonal axes ranked by explained variance, SCA identifies axes with sparse neuronal loadings. These axes are dominated by relatively small groups of strongly weighted (contributing) neurons, allowing the population geometry to be related to specific neuronal subsets. To focus on generalization associated with advance planning, sparse components (SCs) were identified using plan trials, and data from all conditions were subsequently projected onto these axes. A cross-validated grid search returned three SCs in each region (see Methods). Neuronal loadings in both regions were highly sparse (kurtosis values greater than 3 indicate heavy-tailed distributions), arguing that a small subset of neurons contributed disproportionately to the identified geometry. Sparsity was particularly more pronounced in MD (**Fig. 4b**, PL: SC1, *k* = 23.1; SC2, *k* = 19.2; SC3, *k* = 36.1; MD: SC1, *k* = 50.5; SC2, *k* = 25.5; SC3, *k* = 68.7). Together, the SCs accounted for 10.6 % of the total variance of demixed input matrix (i.e., without the condition-independent temporal component) in PL (SC1: 4.7 %; SC2: 3.2 %; SC3: 2.7 %) and 12.9 % in MD (SC1: 4.5%; SC2: 4.3 %; SC3: 4.1 %).

Projection of population activity from all task conditions onto the sparse axes revealed distinct organizations in PL and MD. PL SC1 differentiated left-plan from right-plan trials during the context delay and early instruction epoch (**Fig. 4c**). Dominant SC1 neurons (loading > 2 SD; n = 30; **Fig. 4d**) strongly discriminated left from right conditions in both epochs and showed high cross-epoch generalization (**Fig. 4e, f**, Wilcoxon signed-rank test to 0.5 baseline, *p* < 0.001). PL SC2 and SC3 were both strongly modulated by the defer context but were engaged during different phases of the instruction epoch (**Fig. 4g, k**), consistent with the biphasic PL response observed previously (**Fig. 2c**). Dominant SC2 neurons (n = 27; **Fig. 4h**) showed greater cross-epoch generalization (**Fig. 4i, j**, Wilcoxon rank-sum test, *p* < 0.001) than dominant SC3 neurons (n = 30; **Fig. 4l-n**).

In MD, none of the sparse components showed comparable context modulation (separation between plan and defer trials as for PL SC2 and SC3) during the context epoch (**Fig. 4o-w**). Instead, activity separated left from right trials. Although MD components also discriminated instructed side during the instruction epoch, dominant MD neurons (matched to the PL subset sizes using loading > 1.5 SD; n = 25 for SC1, n = 26 for SC2, and n = 20 for SC3) showed smaller cross-epoch generalization than PL (**Fig. 4p-z**, Wilcoxon rank-sum test between every pair of PL SC1/SC2 to MD SC1/SC2/SC3, *p* < 0.001).

Removing dominant SC neurons abolished above-chance cross-epoch generalization in the remaining PL and MD populations (PL: n = 542; MD: n = 349; **Fig. S4**), indicating that generalization depended disproportionately on these sparse neuronal subpopulations rather than being weakly distributed across many neurons.

In summary, dSCA identified functionally distinct sparse neuronal subspaces in PL and MD. PL contained a sparse component whose instructed-side representation generalized across context and instruction epochs. By contrast, MD sparse components primarily represented instructed side within individual task epochs and showed little cross-epoch generalization.

## Discussion

In this study, we examined how population activity in PL and MD differed when context allowed an upcoming response to be planned versus when the required response remained unspecified until the instruction cue. This task therefore isolates a dimension of cognitive flexibility beyond the typically studied switching between stimulus-response mappings or choosing between simultaneously available options: it allows us to study the processes contributing to the early formation of action representations. Both PL and MD represented response-side information, but they differed in its timing and cross-epoch generalization. The principal dissociation was earlier prospective response-side information in MD and a more strongly generalizable representation across context and instruction epochs in PL.

To organize the temporal results, we refer to the post-context epoch, the early post-instruction epoch and the later post-instruction epoch as planning, initiation and executive stages, respectively. During the planning stage, MD population activity distinguished the prospective instructed side earlier than PL, and context-related response-side information was more prominent than in PL. This profile indicates rapid recruitment of MD population activity when context permits advance planning. Previous studies have found that MD inputs can sustain prefrontal activity during working-memory maintenance (Bolkan et al., 2017), amplify local cortical connectivity associated with maintenance of rule representations (Schmitt et al., 2017) and influence switching between prefrontal cortical representations (Rikhye et al., 2018). Our results extend these findings by suggesting that MD also takes a lead over prefrontal cortex when action representations can be formed in advance without having to wait for additional information.

During the early instruction period, PL, but not MD, displayed a planning-related elevation in neuronal activity, and response-side information generalized from the context epoch into the instruction period more strongly in PL than in MD. PL activity also separated into early and later instruction components at the population subspace (sparse component) level. These properties are consistent with a broader view of prefrontal cortex as carrying prospective, goal-oriented information rather than only stimulus identity or low-level motor variables (Funahashi & Andreau, 2013; Hanganu-Opatz et al., 2023; Rautio et al., 2026; Spellman et al., 2021). High-dimensional, mixed-selective prefrontal populations can represent multiple task variables (Fusi et al., 2016; Parthasarathy et al., 2017), including in partly orthogonal spaces (Rautio et al., 2026; Xie et al., 2022). In our task, SCA identified both task-stage-specific subspaces and a PL component in which response-side information generalized across context and instruction epochs. Thus, representational geometry changed across task stages while retaining a partially shared response-side axis, consistent with related accounts of prefrontal neural geometry that reflects cognitive operations, not cognitive content (Tang et al., 2020; Wang, 2025; Wang et al., 2025).

Recent findings also indicate that PFC can reuse low-dimensional subspaces for shared task information across tasks (Osako et al., 2025; Park et al., 2025; Tafazoli et al., 2026). Our observation of convergence toward a context-invariant instructed-side representation in the executive stage is consistent with reuse of a shared low-dimensional code. Such a representation may provide a format suitable for downstream motor readout (Binish et al., 2026; Shenoy et al., 2013). Notably, stable population-level representations coexisted with dynamic single-neuron activity and changing task-epoch selectivity. This coexistence illustrates the advantage of low-dimensional, populational coding which preserves task-relevant information while enabling flexible recruitment of single neurons (Murray et al., 2017; Spaak et al., 2017; Stokes, 2015).

Although MD also contained multiple subspaces across task stages, its response-side representations showed little cross-epoch generalization. This pattern suggests that, while PL activity was characterized by more abstract, high-dimensional representational formats, MD activity was dominated by simpler, demixed codes for a few specific, immediately task-relevant variables (DeNicola et al., 2020; Lam et al., 2024).

In conclusion, our recordings reveal complementary population dynamics in PL and MD during context-dependent response planning. The findings suggest a division of representational labor within the prefrontal-thalamic system: MD emphasizes the timing of prospective response formation, whereas PL carries a more stable, sparsely implemented representation across successive task states. This organization provides a population-level framework for understanding how advance information is carried forward during flexible, context-sensitive action planning. Because the study was observational, it does not establish causal interactions, information flow or control within the PL-MD circuit. Future work combining circuit dissection with pathway-specific manipulation, ideally in tasks that dissociate cue identity from response side, should test how prefrontal-thalamic interactions mechanistically shape the timing and stability of action representations and thereby support cognitive flexibility.

## Methods

### Animal procedures

All procedures were conducted in accordance with protocols approved by the local government authority (Regierung von Oberbayern). Five male wildtype C57BL/6 mice were used for behavioral training and electrophysiological recordings, and a separate cohort of five mice was used for anatomical tracing experiments. For the main experiment, animals underwent surgery for headbar implantation between 8 and 10 weeks of age. After surgery, they were housed individually with ad libitum access to food and water under a reversed 12 h/12 h dark-light cycle. During behavioral training and electrophysiological recording, animals were placed under a water-restriction protocol and received water rewards for correct responses. Daily water intake was maintained at a minimum of 1500 µL throughout training. Once animals reached the behavioral criterion, a second surgery was performed to prepare craniotomies for acute recording.

### Surgery

Animals were anesthetized with isoflurane (2 % vol/vol in O₂), and body temperature was maintained at 37.5 °C using a heating pad and monitored continuously with a rectal probe. Preoperative analgesia (metamizole, 200 mg/kg body weight) was administered subcutaneously. Animals were placed in a stereotaxic frame (Model 900LS, Kopf Instruments, USA) using ear bars, and anesthesia was maintained with 0.8-1.5 % isoflurane. Eyes were protected with ophthalmic ointment (Augen- und Nasensalbe, Bepanthen, Germany). A local anesthetic (2 % lidocaine in 0.9 % NaCl) was injected subcutaneously at the incision site. The scalp was disinfected with 70 % ethanol, and the skull was exposed. Periosteal tissue was removed, and the skull surface was roughened using a scalpel blade to improve adhesion. A small craniotomy was made posterior and lateral to lambda for placement of a stainless-steel ground screw using a dental drill (Neurostar, Germany). A thin layer of light-curable adhesive (Optibond All-in-One, Kerr, Germany) was applied to the skull.

Entry sites for later acute recordings were stereotaxically marked by drilling into the cured adhesive and superficial skull. An aluminum headbar was aligned and fixed to the skull with dental cement (Tetric EvoFlow, *Ivoclar Vivadent, Germany*). The remaining exposed skull was covered with multiple layers of dental cement to ensure mechanical stability, while the marked recording sites were sealed with silicone elastomer (Kwik-Cast, *World Precision Instruments, UK*).

Animals were given ad libitum access to water on the day before craniotomy surgery. Presurgical procedures were performed as described above. The silicone covering the recording sites was removed, and the skull surface was cleaned with 70 % ethanol followed by 0.9 % NaCl. Craniotomies were opened at the stereotaxic marks made during the initial surgery and then resealed with silicone elastomer. Post-operative analgesia (meloxicam, 1.5 mg/kg body weight) was administered subcutaneously immediately after surgery and once daily for three days thereafter.

### Behavioral setup

Behavioral training and recording took place in a custom-built sound-attenuating chamber. Mice were head-fixed with their bodies supported in a plastic tube, allowing free movement of their forelimbs. The forepaws rested on a spherical response device (“response ball”). Water was delivered via a blunt cannula placed near the mouth and dispensed by an Arduino-controlled syringe pump (NE-500, *New Era Pump Systems, USA*).

Visual stimuli were presented on a 10-inch LCD monitor (FT10TMB, *Faytech, Germany*) positioned 15 cm in front of the animal. Auditory stimuli were delivered through a pair of electrostatic speakers (ES1, *Tucker Davis Technologies, USA*), driven by electrostatic amplifiers (ED1, *Tucker Davis Technologies, USA*).

Left and right responses were measured using a ping-pong ball mounted on a metal rod, allowing only horizontal (left-right) rotation. A diametrically magnetized disc magnet attached to the rod’s anterior end enabled angular displacement tracking via a Hall-effect rotary encoder (MIB22H, *Megatron, Germany*). The encoder output was converted into a continuous analog signal through a custom circuit (designed by Christian Obermayer), then fed into a breakout panel (BNC-2090A, *National Instruments, USA*) connected to a data acquisition system (PCIe-6321, *National Instruments, USA*). TTL pulses were sent to reset the analog signal.

Behavioral task control was implemented using custom MATLAB scripts (*MathWorks, USA*), built on MonkeyLogic 1 (Asaad et al., 2013) and the NIMH Toolbox (Hwang et al., 2019).

### Task and stimuli

Each trial began with the presentation of a grey screen (0.1 luminance). At the same time, the virtual position of the response ball was reset to baseline, and the animal was required to hold the ball still within ±2.5 mm for 500 ms (initiation epoch). If this threshold was exceeded, a TTL pulse was sent to reset the ball signal and extend the initiation period by an additional 500 ms as temporal punishment. If the ball remained within the threshold for the required duration, one of three auditory context cues (61 dB SPL at the mouse’s head position) was played binaurally for 100 ms.

In 25 % of trials, a low-pass filtered white noise (< 8 kHz) signaled the left-plan context, which was always followed by a go-left instruction. In 50 % of trials, unfiltered white noise indicated the defer context, which was followed randomly by either a go-left or go-right instruction (50 % each). In the remaining 25 % of trials, a high-pass filtered white noise (> 14 kHz) signaled the right-plan context, which was always followed by a go-right instruction. Following the context cue, the ball signal was reset again, and the animal was required to keep the ball still for another 500 ms (delay epoch). As in the initiation epoch, any movement exceeding ±2.5 mm reset the signal and extended the delay.

After the delay, one of two movement instruction cues (78 dB) was presented for up to 1000 ms: a sinusoidal up-sweep (11 kHz to 14 kHz) served as the go-left instruction, whereas a sinusoidal down-sweep (15 kHz to 12 kHz) served as the go-right instruction. Animals were allowed to respond by rotating the ball left or right for up to 5000 ms after instruction onset. A response was detected when the ball moved beyond ±8 mm; if the instruction sound was still playing, it was then terminated. Correct responses were rewarded with a water drop delivered over 500 ms. The grey screen was then turned off, followed by a 5000 ms intertrial interval. To train animals to perform the full task, two preliminary stages were introduced:

#### Movement Instruction Stage

After habituation to the head-fixed setup, animals were initially trained using only *go-left* and *go-right* instructions. They were allowed up to 20 s to move the wheel beyond a small threshold. Initially, instructions were presented in blocks of 20 to 30 trials. As proficiency increased, block sizes were reduced (eventually to single trials), response thresholds were increased (from ±3.5 mm to ±8 mm), and reward size was gradually decreased (from 8 µL to 4 µL) to encourage sustained engagement.

#### Context Integration Stage

Once animals reached >90 % performance across consecutive sessions, context cues and the delay period were introduced. Animals were trained to withhold movement during the delay. The delay interval gradually increased from 50 ms to 500 ms as proficiency improved.

### Trial screening and psychometrics

To exclude periods of fluctuating motivation, warm-up and cool-down phases of each session were discarded using the following procedure. Contiguous blocks of trials were labeled “bad runs” if the running-average performance over 15 trials fell below 50 %. The first non-bad run after the initial 20 trials was designated as the session start for valid trials. Similarly, the session endpoint was defined from continuous sequences of miss or incorrect trials. All miss trials (i.e., no response detected within the permitted response window) were excluded from the analysis. A session was considered valid if it included at least 100 valid trials, at least 70 % overall performance, at least 50 % performance in each condition, and at least 5 correct trials per condition.

For neuronal data analysis, only sessions with histologically verified silicon probe placements were included. Accordingly, one animal’s dataset was excluded from analysis. Session performance was defined as the fraction of correct responses among valid trials. Response time was defined as the interval from instruction onset to the moment the response threshold was crossed. The effects of context and instruction on behavior were evaluated using repeated-measures ANOVA with context (plan vs. defer) and instruction (go-left vs. go-right) as factors. No violations of the assumption of equal variance were detected using Levene’s test.

Behavioral metrics were visualized using Raincloud plots, which combine scatter plots, boxplots, and violin plots to simultaneously show individual data points and distribution features. In these plots, each dot represents the session average of a behavioral metric; half-violins show kernel density estimates (KDE); boxplots show interquartile ranges with medians indicated by white dots; and whiskers denote 1.5 times inter-quartile range (IQR).

### Submovements decomposition

To analyze movement microstructure, we extracted minimum-jerk submovements (SMs) from the response ball velocity, calculated as the temporal derivative of the analog voltage signal tracking wheel position. Periodic TTL reset pulses (lasting ∼33 ms) were used during the experiment to reset the voltage signal back to 0 V, introducing discontinuities into the trace. These discontinuities were interpolated using Gaussian Process (GP) regression with a two-component kernel, implemented via scikit-learn (https://github.com/scikit-learn/scikit-learn).

The first kernel component was a Radial Basis Function (RBF), which weighed nearby data points more heavily than distant ones. Given that reset pulses lasted up to 33 ms and were typically flanked by ≥2 data points, the RBF length scale ℓ was fixed at 37 ms (Duvenaud, 2014). The second kernel component was a white noise kernel to model measurement noise, with variance σ² determined per recording setup.

Following interpolation, velocity traces were fitted with a linear combination of bell-shaped polynomial basis functions:

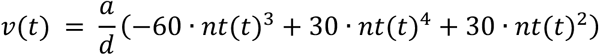

where *v*(*t*) is the velocity, *a* is the movement path length, *d* is the duration, and *nt*(*t*) is the normalized time 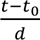; *t*_0_ is the time of movement onset.

The fitting procedure comprised three iterative steps: proposal, fitting, and evaluation. During proposal, the residual from the current best-fitting model was used to identify a candidate submovement. In the first iteration, the model was initialized to zero, so the residual equaled the velocity trace. We first extracted the local maxima and their neighboring local minima of the velocity residual (*SciPy*, https://github.com/scipy/scipy/tree/v1.5.3). The temporal distance between the neighboring local minima was used as the initial estimate of submovement duration. The proposed duration *d* and velocity peak *v_peak_* were used to calculate the corresponding path length *a* of each submovement:

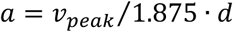

The proposed SM with the longest path length was advanced to the fitting stage if it was not entirely nested within another submovement and had not been included in a previous iteration. During fitting, we used nonlinear least-squares optimization (*lmfit*, https://lmfit.github.io/lmfit-py/) to find the optimal *a*, *d*, and *t*_0_ of the proposed SM. The model was evaluated after each iteration. If the Bayesian information criterion (BIC) improved relative to the preceding iteration, the fitted SM was accepted. Otherwise, the current SM was rejected, and the fitting procedure was repeated for the next-longest proposed SM.

Further analyses of SM count and amplitude focused on two time windows: (1) Subthreshold movement, during the context and delay period (0–600 ms after context onset), and (2) Response movement, from instruction onset to response threshold crossing. Reconstructed direction-specific velocity traces were derived by summing SMs with leftward or rightward movement direction during each epoch. A side-index (SI) was computed to capture lateral movement bias per trial:

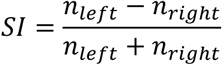

Movement onset time was defined as the start of the first fitted SM after instruction onset. Movement execution time was defined as the interval from movement onset to response-threshold crossing. Repeated-measures ANOVA was used to assess context and context × instruction effects. Pairwise condition comparisons were performed using Wilcoxon signed-rank tests.

Trials were further categorized by movement tendency (leftish vs. rightish) based on cumulative path length during the first 600 ms after context onset. Tendency was computed as the normalized signed sum of path lengths. Trial counts were balanced between decoder categories for further analysis.

### Retrograde tracing

To trace MD-to-PL projections, 75 nL of cholera toxin subunit B (CTB) conjugated to different fluorophores (CTB 488, CTB 555, or CTB 647) was injected into the anterior (aPL), middle (mPL), or posterior (pPL) regions of PL in adult mice (n = 3 per group). Brain sections were collected and imaged to quantify retrogradely labeled neurons in MD, which was segmented into medial (MDm), central (MDc), and lateral (MDl) subdivisions. The anterior-posterior distribution of MD projection neurons was analyzed to characterize connectivity patterns.

To trace PL-to-MD projections, CTB 488 or CTB 647 was injected into the medial (MDm) or lateral (MDl) subdivisions of MD, respectively (*n* = 2 for MDm, *n* = 3 for MDl). Retrogradely labeled neurons were quantified in medial prefrontal cortex regions, including secondary motor cortex (M2), dorsal cingulate cortex (Cg1), prelimbic cortex (PL), and infralimbic cortex (IL). Confocal microscopy was used to acquire images at both 4× and 20× magnifications to confirm tracer localization and labeling in targeted regions.

### Electrophysiology

Before acute recording, animals were head-fixed in the behavioral setup. A custom-made reference cable connected the implanted skull screw to the reference electrode of the silicon-probe adapter. The silicone elastomer covering the craniotomies was removed, and the craniotomies and brain surface were rinsed with 0.9 % saline. One silicon probe per site (A1x32-Poly2-10mm-50s-177-A32 for dual-site recordings and A2x32-Poly2-6mm-23s-160 for single-site recordings; NeuroNexus Technologies, USA) was positioned at the brain surface using a micromanipulator (Luigs-Neumann, Germany). With depth referenced to the brain surface, the probes were slowly lowered to the target. The well surrounding each craniotomy was filled with 2 % agarose in 0.9 % saline to improve stability and prevent drying of the brain surface. After insertion, probes were allowed to settle for 10-30 min. Recording began shortly before the behavioral session. After the session, probes were retracted, the craniotomies were rinsed with 0.9 % saline, and the openings were covered with silicone elastomer. Recordings were obtained from left and right hemispheres in different animals.

Data acquisition was performed using the OmniPlex Neural Recording Data Acquisition System (Plexon, USA). The silicon probes were mounted onto a probe adaptor (Connect HST/32V, Plexon, USA), then to an analog headstage (HST/32o25-GEN3-36P-G1, Plexon, USA). The headstage was connected through a digitizing amplifier (DigiAmp, Plexon, USA) to the OmniPlex chassis, which then passed the signal to the recording PC via the OmniPlex software. Signal was sampled at a rate of 40 kHz. The wideband signal was split into field potentials (low-pass filtered at 500 Hz and downsampled to 1000 Hz) and spike signals (high-pass filtered at 300 Hz).

### Spike sorting and unit selection

The continuous spike signals were fed to Kilosort 1 (Pachitariu et al., 2016) for automatic spike sorting based on the spatiotemporal features of spike waveforms. Kilosort utilizes template matching and batch-based optimization to efficiently handle large-scale electrophysiological datasets. Curation of the pre-clustered data (up to 512 clusters per session) was done with Phy (https://github.com/cortex-lab/phy). Clusters were merged if their spatiotemporal waveforms or autocorrelation histograms were very similar. Drifts in amplitude or spatial location (towards neighboring channels) may split the recording of a single neuron into separated clusters; this was also manually curated. Good single-units were chosen conservatively based on these criteria: (1) distinct, often large amplitude waveforms; (2) physiologically plausible waveform shapes; (3) a physiologically plausible refractory period. If there were clear violations of refractory period but a physiologically plausible waveform, the cluster was labelled as MUA. Splitting was not performed. Finally, clearly non-physiological waveforms were labelled as noise.

For subsequent analyses, we excluded periods of sparse firing at the beginning and end of each unit’s recorded lifetime to improve firing-rate stationarity. This was necessary because units sometimes drifted into or out of the recording during acute sessions. Only activity considered to originate from single units was included. Single units were further screened for an average firing rate of at least 1 Hz and presence in at least 5 correct trials per condition. A unit was defined as present if it showed no more than 5 consecutive trials with a low firing rate (< 1 Hz).

### Standardized firing rate

Binary spike trains were first downsampled to 1000 Hz and then smoothed into continuous firing-rate time series by convolving with a Gaussian window of σ = 25 ms and length of n = 9 σ. Each neuron’s time series *fr*(*t*) were standardized to baseline by:

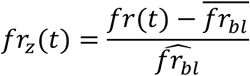

where 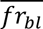 is the mean baseline firing rate and 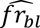 is its standard deviation. The baseline period was defined as −200 to 0 ms relative to trial onset.

The change in firing rate was estimated as the first temporal derivative, calculated as the difference between consecutive samples (1 ms resolution), resulting in a dFR/dt expressed in units of z/ms.

### Zeta-test derived metrics

Neuronal responsiveness to stimuli was computed from the raw spike timestamps using the parameter-free ZETA (Zenith of Event-based Time-locked Anomalies) test (Montijn et al., 2021). ZETA-test computes the magnitude of responsiveness (ζ) and a ζ-derived instantaneous firing rate without binning the data and is therefore independent of the arbitrary choice of analysis parameters. To achieve this, single-unit spikes were first aligned to the stimulus onset with a systematically varied offset *τ*. The empirical cumulative distribution of the observed spike trains *F_obs_*(*τ*) within the stimulus interval [*t*_start_, *t*_end_] was compared to a null distribution generated by averaging across 100 resamples of the spike trains. The most extreme deviation across different offsets *τ* was taken as the ζ value of the unit, and the time of peak ζ-derived firing rate was taken as its response latency:

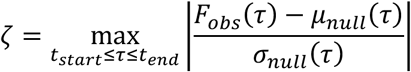

where *μ_null_*(*τ*) and *σ_null_*(*τ*) denote the mean and standard deviation of the null distributions. A single unit was deemed responsive to a stimulus if its ζ value (responsiveness magnitude) exceeded 1.96 (i.e., *α* = 0.05, two-tailed). Preference was defined as the stimulus to which a neuron’s activity was maximally modulated.

Neuronal recruitment in each region was approximated from the distribution of single-unit latencies. Unimodality of each latency distribution was assessed using Hartigan’s dip test. Regional response timing was summarized by the first mode of a kernel density estimate (KDE), corresponding to the most frequent value in the estimated distribution. Relative timing between PL and MD was assessed by comparing these modes using nonparametric bootstrap resampling.

The association between single-unit cue preferences in the context and instruction epochs was evaluated using a chi-square (*χ*^2^) contingency test. The *χ*^2^ statistic compared the observed joint distribution with the distribution expected if cue preferences in the two epochs were unrelated.

### Neuronal information

To quantify firing-rate information about instructed response side (left/right), we calculated the percentage of explained variance (ω^2^ PEV) (Buschman et al., 2011) using:

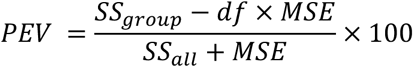

where *df* denotes the degrees of freedom, *MSE* denotes the mean squared error, and *SS* denotes the sum of squares obtained from the ANOVA. PEV was calculated separately for plan and defer trials.

### Granger-causality analysis of spike trains

As an exploratory measure of condition-dependent directed statistical dependence within each recorded region, we computed Granger-causality (GC) values from binary spike trains of sorted single neurons (Granger, 1969). We used an unconditional, nonparametric Granger-Geweke approach for point processes (Geweke, 1982), implemented in custom Python scripts adapted from MATLAB code (Dhamala et al., 2008). GC was evaluated for neuron pairs within PL or within MD; it was not used to measure PL-to-MD or MD-to-PL communication. Because the analysis is observational and pairwise, GC is interpreted here as directed predictability rather than proof of a causal or anatomical connection. Multitaper spectral-density estimation (Bokil et al., 2010) was applied to binned spike data to obtain the mean cross-spectrum across trials. Relevant channels and time bins were selected to improve stationarity, a set of Slepian tapers (DPSS) was generated, and each taper was multiplied by the binned data for each channel. A fast Fourier transform (FFT) was then applied to the tapered signals. For each pair of channels, the product of one channel’s complex spectrum and the conjugate of the other channel’s spectrum was averaged across tapers and trials to estimate cross-spectral density. Mathematically, the cross-spectral density between channels i and j at frequency f was expressed as:

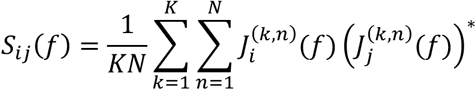

where 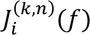 is the Fourier transform of the data from neuron i, tapered by the k-th taper in the n-th trial, * denotes the complex conjugate, *K is the number of tapers, and N* is the number of trials.

We then used Wilson’s spectral-factorization algorithm to decompose the two-neuron spectral-density matrix S(ω) into a transfer function H(ω) and an error-covariance matrix Z. First, we performed an inverse fast Fourier transform (IFFT) and obtained an initial estimate for the factorization. In each iteration of the algorithm, we refined this estimate by minimizing the difference between the reconstructed spectral density and the original spectrum until the change in the error fell below a specified tolerance. At each frequency ω, the final spectral density was expressed in the factorized form:

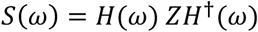

where *H*^†^(*ω*) denotes the Hermitian transpose of *H*(*ω*), and *Z* is the constant (frequency-independent) error covariance matrix. This procedure yielded the factorized spectral density, transfer function, and error- covariance matrix. The frequency-dependent Granger-Geweke measure from neuron i to neuron j was then calculated by comparing the modeled power of neuron j with the contribution associated with neuron i included versus excluded:

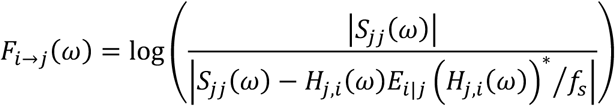

where *S*_jj_(*ω*) is the factorized power spectrum of neuron j, *H*_j,*i*_(*ω*) is the transfer function from neuron i to neuron j, * denotes the complex conjugate, *E_i_*_|j_ is the corrected covariance, and *f_s_* is the sampling frequency.

A frequency-integrated Granger-Geweke value was obtained by integrating across frequencies *ω* ∈ [0, *π*]:

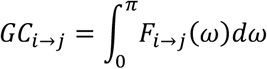

For each neuron pair, significance of the GC value was assessed against a null distribution generated from 200 trial shuffles using a per-pair threshold of p < 0.05.

### Population decoding: support vector machine (SVM)

To assess population coding of task variables, we implemented L2-regularized linear support vector machines (scikit-learn). SVMs for all recorded neurons or selected dominant-neuron subsets in PL and MD were trained with hinge-loss, stopping tolerance *tol* = 1 × 10^−5^, regularization *C* = 100, and maximum number of iterations *niter* = 10000. For the much smaller dominant-neuron subsets, SVMs were trained with hinge-loss, stopping tolerance *tol* = 1 × 10^−3^, regularization *C* = 1, and maximum number of iterations *niter* = 10000. Binary spike trains were smoothed by convolution with a Gaussian window (*σ* = 25 ms) and downsampled to 40 Hz. Recorded neurons were pooled across all valid sessions into region-specific pseudopopulations (Meyers, 2018). For each neuron, 50 trials were sampled with replacement and concatenated across neurons. This procedure removes trial-by-trial neuronal covariation; decoding therefore reflects information in the sampled single-neuron activity distributions rather than information carried by simultaneous covariation. The training fraction was 0.75, corresponding to a 3:1 training-to-test ratio. This procedure was repeated for 100 shuffles. Neural information was quantified as the mean accuracy of class predictions of trained classifiers on held-out test trials. Means and standard deviations were calculated across pseudotrial shuffles.

Cross-temporal decoding was performed by training a classifier on activity from one time bin and testing it on activity from another. Two approaches were used to assess context-dependent changes in response-side coding during the instruction epoch. First, each fitted linear SVM defined a multidimensional hyperplane that separated the two classes. The cosine similarity *cos*(*θ*) between the weight vectors of two classifier hyperplanes 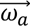 and 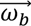 was calculated via:

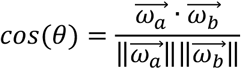

The centered dot denotes the dot product. Cosine similarity was calculated for every pair of pseudotrial shuffles and then averaged. Second, cross-context decoding was performed analogously to cross-temporal decoding. Classifiers were trained on a subset of trials in one context and tested in another context. Cross-context accuracy was compared with within-context accuracy.·

The generalization index (GI) was defined as the mean cross-epoch decoding accuracy obtained by training the decoder on context-epoch activity and testing it on instruction-epoch activity within the first 200 ms of each epoch.

### Demixing sparse component analysis (dSCA)

To identify sparse components in population activity, we implemented sparse component analysis (Lin et al., 2023), which reduced data dimensionality while penalizing nonsparse neuronal loadings onto each component.

To construct the pseudopopulation activity matrix, the normalized firing-rate vector of each neuron was first demixed by subtracting the overall mean and mean temporal fluctuation from that of each analyzed condition, yielding demixed plan-condition vectors for left-plan and right-plan conditions, respectively.

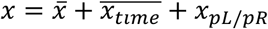

We then reshaped the activity matrix by concatenating across the two plan conditions and two epochs: (0 – 600 ms from context onset and 0 – 600 ms from instruction onset) into a two-dimensional matrix *X* ∈ ℝ*^P^*^×*N*^, where *P* = *n_conditions_* × *T*, and *N* denotes the number of neurons. The resulting activity matrix X was approximated by the sum of k component-activity vectors 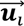 and their corresponding neuronal loading vectors 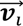 :

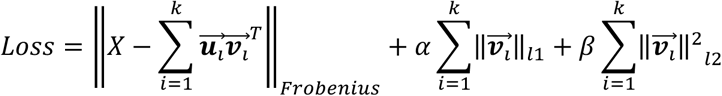

subject to the constraint 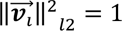. The columns of X comprise *n*_units_ neuronal features, and the rows comprise *n*_conditions_ × *n*_time_condition-time observations. Hyperparameter *α* controls the strength of L1 regularization, which promotes sparsity of the neuronal loadings. Hyperparameter *β* controls the strength of L2 regularization, which stabilizes the loss landscape across random initializations. Hyperparameters *α*, *β*, and *k* were determined with a twofold cross-validated grid search.

This procedure yielded the following matrix factorization:

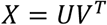

where *U* contains component responses across condition-concatenated time, and V contains neuronal loading vectors for the sparse components. Defer trials were subsequently incorporated by projecting a full-condition activity matrix *X_full_* containing demixed activity from all four conditions (Lp, Rp, Ld, and Rd) using the same loading matrix. Component responses were obtained by the following linear least-squares solution:

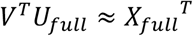

## Data Availability

Data are available on request from the corresponding author.

## Code Availability

Custom code for behavior analysis and neuronal analysis is publicly available at https://github.com/wangxuanyu0419/JacobLab_MouseDecision.

## Acknowledgements

S.N.J. acknowledges funding for this work from the European Research Council (Starting Grant 758032, MEMCIRCUIT) and the German Research Foundation (DFG) (JA 1999/3-1).

## Author Contributions

D.H., A.R. and S.N.J. conceived the study and designed the experiments. D.H. and A.R. collected the data with assistance from T.W.B. D.H. and A.R. performed the initial analyses. X.W. extended the analyses and contributed the sparse-component analysis. S.A.F. developed the scripts for spike-train Granger-causality analysis under the supervision of X.W. D.H. and X.W. prepared the figures. X.W. and S.N.J. wrote the manuscript with input from D.H. All authors reviewed and approved the final manuscript. S.N.J. supervised the study and provided funding.

---

**Fig. S1.**
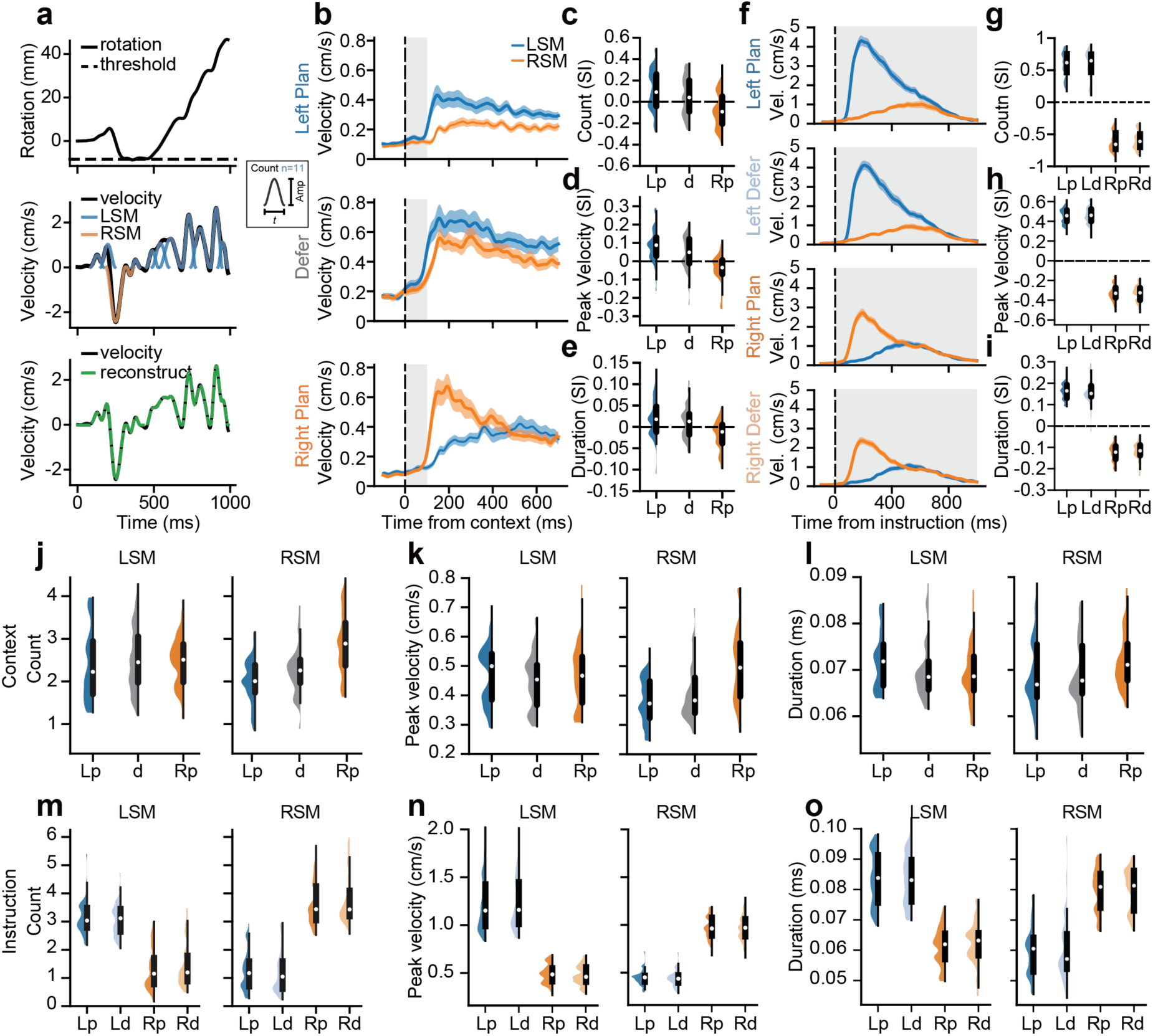
Submovement metrics in the context and instruction epochs. **a.** Illustration of minimum-jerk submovement (SM) fitting. Top, rotation trace from one example trial. Positive and negative values indicate leftward and rightward movements; the dashed line marks the response threshold (10° to the right in this example). Middle, ball velocity (black) and fitted minimum-jerk SMs (colored bell-shaped curves). Inset, fitted SM parameters. Bottom, ball velocity (black) and velocity reconstructed from fitted SMs (green dashed line). **b.** Reconstructed LSM and RSM velocity during the context epoch for the three context cues, averaged across included sessions (n = 55). Solid lines indicate the mean, shaded bands the ± SEM. **c.** Side index (SI) for SM count during the context epoch for the three context cues, calculated for each trial as the difference between LSM and RSM counts divided by their sum. Distribution across sessions are shown as half-violins with box plots indicating the median (white dot), quartiles (thick line), and 1.5 times the interquartile range (thin line). **d.** Same as **c** for SM peak velocity. **e.** Same as **c** for SM duration. **f-i.** Same as **b-e** for the instruction epoch. **j.** LSM and RSM counts in the context epoch. **k.** Same as **j** for SM peak velocity. **l.** Same as **j** for SM duration. **m-o.** Same as **j-l** for the instruction epoch.

**Fig. S2.**
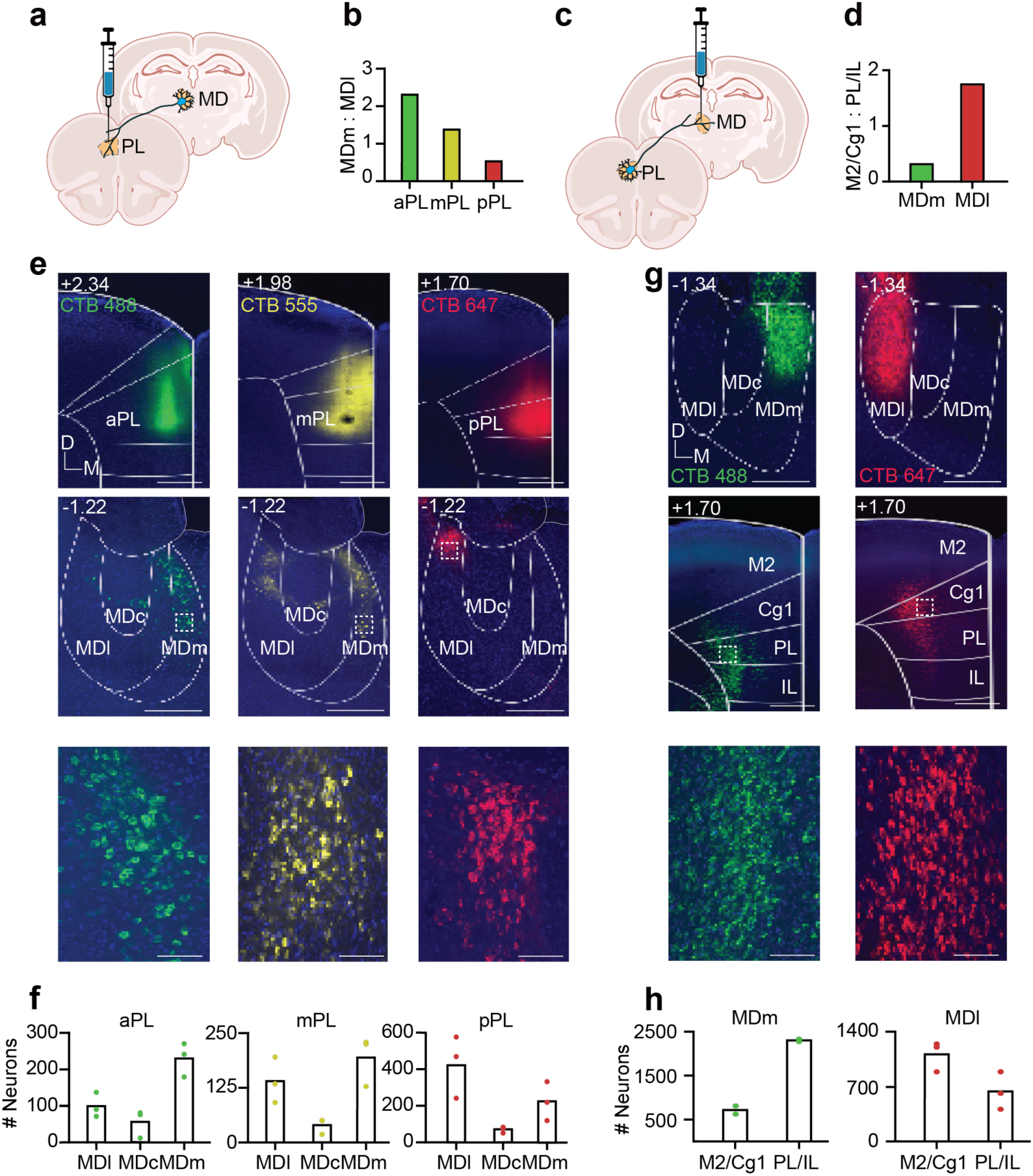
Anatomical tracing of reciprocal projections between PL and MD. **a.** Schematic of retrograde tracing of MD projections to PL (created with BioRender). CTB was injected into PL, taken up by axon terminals, and transported retrogradely to projection-neuron cell bodies in MD. **b.** Distribution of retrogradely labeled neurons across medial (MDm) and lateral (MDl) MD after injections into anterior (aPL), middle (mPL) or posterior (pPL) PL. **c.** Retrograde tracing of PL projections to MD. **d.** Distribution of labeled neurons across secondary motor/dorsal cingulate cortex (M2/Cg1) and prelimbic/infralimbic cortex (PL/IL) after injections into MDm or MDl. **e.** Example confocal images from MD-to-PL tracing. Top, PL injection sites along the anterior-posterior axis (aPL, CTB 488, n = 3; mPL, CTB 555, n = 3; pPL, CTB 647, n = 3); anterior-posterior coordinates are shown at the upper left. Middle, retrogradely labeled neurons in MD, divided into lateral (MDl), central (MDc), and medial (MDm) subdivisions. Bottom, magnified views of the areas indicated by dashed boxes. **f.** Numbers of labeled neurons in MDl, MDc, and MDm after injections into aPL (left), mPL (middle), or pPL (right). Colored dots show individual animals and bars show group means. **g.** Example confocal images from PL-to-MD tracing. Top, MD injection sites (MDm, CTB 488, n = 2; MDl, CTB 647, n = 3). Middle, retrogradely labeled neurons in medial prefrontal regions, including M2/Cg1 and PL/IL. **h.** Numbers of labeled neurons in M2/Cg1 and PL/IL after injections into MDm (left) or MDl (right).

**Fig. S3.**
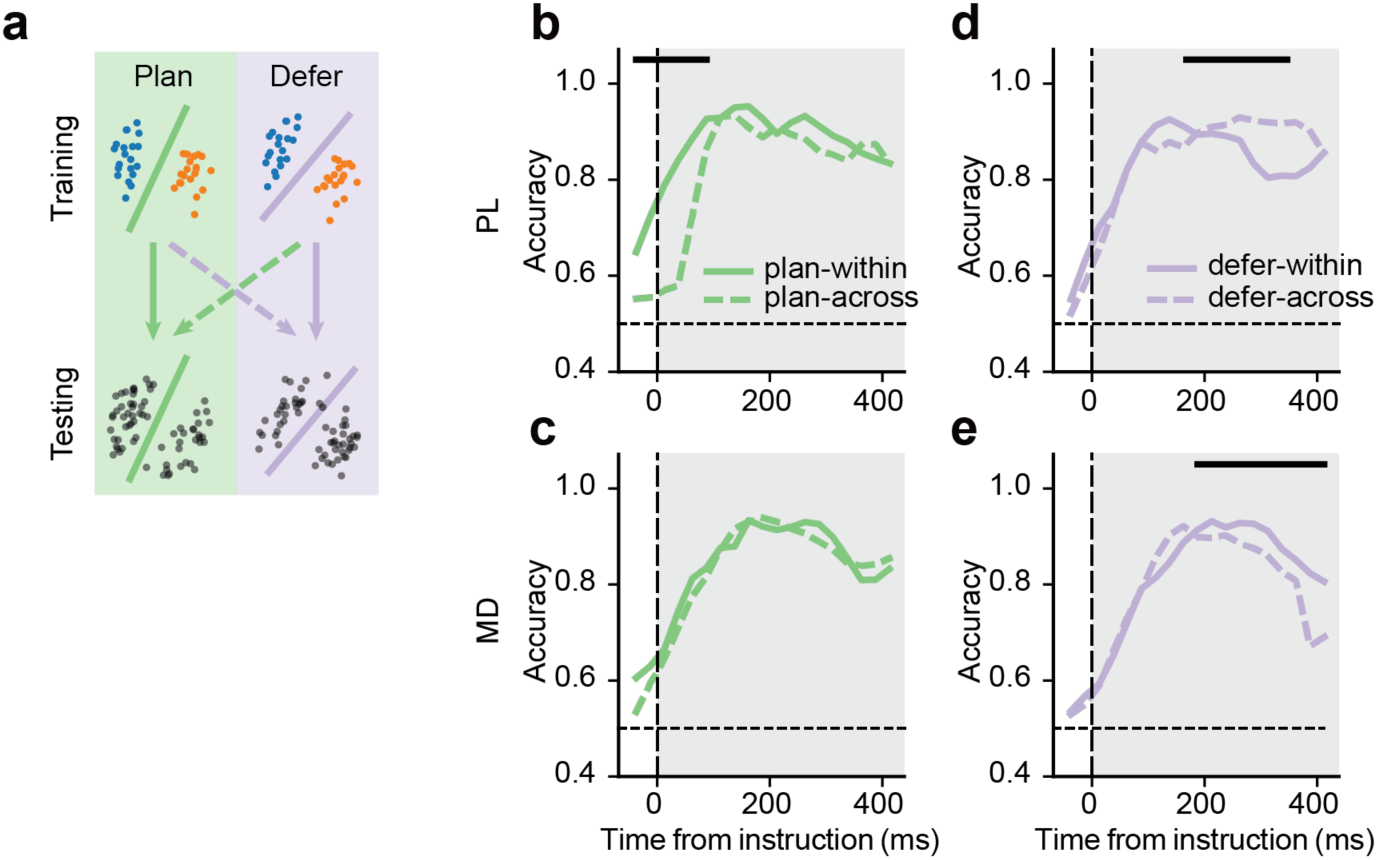
Cross-context decoding in the instruction epoch. **a.** Schematic of within-context and cross-context decoding (solid and dashed lines, respectively) for instructed response side (left vs. right). **b.** Decoding accuracy for the PL population when testing on plan trials. Classifiers were trained either on plan trials (plan-within, solid line) or defer trials (plan-across, dashed line). Horizontal bars indicate significant time windows from permutation tests: thick bars, *p* < 0.01; thin bars, *p* < 0.05. **c.** Same as **b** for MD. **d.** Decoding accuracy for the PL population when testing on defer trials. Classifiers were trained either on defer trials (defer-within, solid line) or plan trials (defer-across, dashed line). **e.** Same as **d** for MD.

**Fig. S4.**
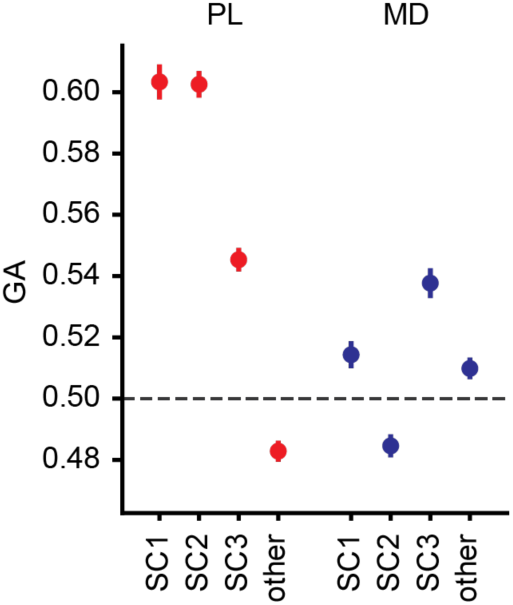
Cross-epoch generalization in sparse-component subpopulations. Cross-epoch decoding accuracy for instructed-side representations (train in context epoch, test in instruction epoch using the first 200 ms of each epoch; generalization accuracy, GA) for neuronal subpopulations composed of dominant neurons with high loadings on sparse components (SC1-SC3) and for non-dominant neurons (other). The horizontal dashed line indicates chance level. Error bars indicate SEM across 100 shuffles.

